# Design and Optimization Protocol for High-Dimensional Immunophenotyping Assays using Spectral Flow Cytometry

**DOI:** 10.1101/784884

**Authors:** L Ferrer-Font, C Pellefigues, JU Mayer, S Small, MC Jaimes, KM Price

## Abstract

Technological advances in fluorescence flow cytometry and an ever-expanding understanding of the complexity of the immune system has led to the development of large 20+ flow cytometry panels. Yet, as panel complexity and size increases, so does the difficulty involved in designing a high-quality panel, accessing the instrumentation capable of accommodating large numbers of parameters, and in analysing such high-dimensional data.

A recent advancement is spectral flow cytometry, which in contrast to conventional flow cytometry distinguishes the full emission spectrum of each fluorochrome across all lasers, rather than identifying only the peak of emission. Fluorochromes with a similar emission maximum but distinct off-peak signatures can therefore be accommodated within the same flow cytometry panel, allowing greater flexibility in terms of panel design and fluorophore detection.

Here, we highlight the specific characteristics regarding spectral flow cytometry and aim to guide users through the process of building, designing and optimising high-dimensional spectral flow cytometry panels using a comprehensive step-by-step protocol. Special considerations are also given for using highly-overlapping dyes and a logical selection process an optimal marker-fluorophore assignment is provided.

## INTRODUCTION

Flow cytometry has rapidly evolved over the past few decades after being first introduced as a one laser system capable of measuring one fluorescence parameter in the late 1960s^1^. In 1979, an improved dual laser multi-parameter flow cytometer was developed that could measure, quantify and sort mammalian cells^2^. Thereafter, multi-laser flow cytometers developed at a rapid pace with three lasers and 10 parameters by the late 1990s^3^, four lasers and 19 parameters by 2004^4^ and up to ten-lasers recently, which has allowed the successful detection of 28 fluorescent parameters^5^.

In parallel, the development of spectral flow cytometry began in 1979^6^, with many different prototypes being trialled^7^ before a commercial instrument was released by Sony in 2012^8^. Whereas conventional flow cytometers record portions of the light spectrum specific for the peak emission of each fluorophore using relevant optical filters, spectral cytometers record small segments of light across the full emission pattern of each molecule^8,9^. Thus dyes not designed for standard optical configurations, or fluorochromes that have identical peaks of emission but different off-peak spectra, can be efficiently differentiated using the unmixing algorithms of spectral cytometry^10^.

Most flow cytometers measure incident photons using Photomultiplier Tubes (PMTs), which are costly, high gain detectors whose quantum efficiency steeply declines after 650nm. The recent use of Avalanche Photodiodes (APDs), which have a high quantum efficiency ranging from 400 to 1100 nm, markedly improves performance in these red and near infrared ranges^11^. In 2017, Cytek Biosciences released the Aurora spectral flow cytometer, which combines spectral flow cytometry with improved detection using APDs. The base unit from Cytek offers 3 lasers (405nm, 488nm and 640nm) and 48 fluorescence channels that use dispersive optics to distribute the collected light across a large detector array, thus allowing the full spectrum from each particle to be measured^7^. Additional laser lines, such as the 561nm yellow-green laser (released in 2018) and the 355nm UV laser (released in 2019), currently give this system 64 fluorescent channels, which allows for the detection of more than 30 commercially-available fluorophores. Previous constraints on the design of high-dimensional flow cytometry panels, which arose from the limited ability to combine and separately detect different fluorochromes, have therefore been replaced by limitations on the availability of the number of dyes with distinct fluorescent spectra^12^.

While spectral flow cytometry has significantly increased the flexibilities of fluorochrome selection and detection, considerations around panel design from conventional flow cytometry still apply. They include a prior knowledge of the biology of the assay, the instrumentation, the expression levels of the markers of interest, the brightness of the selected fluorochromes, and a careful optimization of the antibody panel. This protocol summarizes these points and outlines a number of specific considerations for spectral flow cytometry. It furthermore features a step-by-step guide for the successful design and optimization of a high-dimensional panel for a three-laser Aurora spectral flow cytometer, which can easily be adapted to any spectral flow cytometer with other specifications.

High-dimensional spectral flow cytometry panels are then further analysed using high-dimensional data analysis algorithms (e.g. tSNE, SPADE, FlowSOM, etc.^13^). However, high-quality data are a requirement, as panels with significant spreading errors can result in artificial populations, as well as miss unresolved populations^14^.

## STEP-BY-STEP PROTOCOL TO DESIGN HIGH-DIMENSIONAL PANELS FOR SPECTRAL FLOW CYTOMETRY

In this section, we provide a simple step-by-step protocol for the successful design of a high-dimensional spectral flow cytometry panel. While the design of spectral flow cytometry panels follows similar steps to those described for conventional flow cytometry^5,15^, important differences and additional considerations apply for spectral flow cytometry, which are described in this protocol.

**Figure 1.**
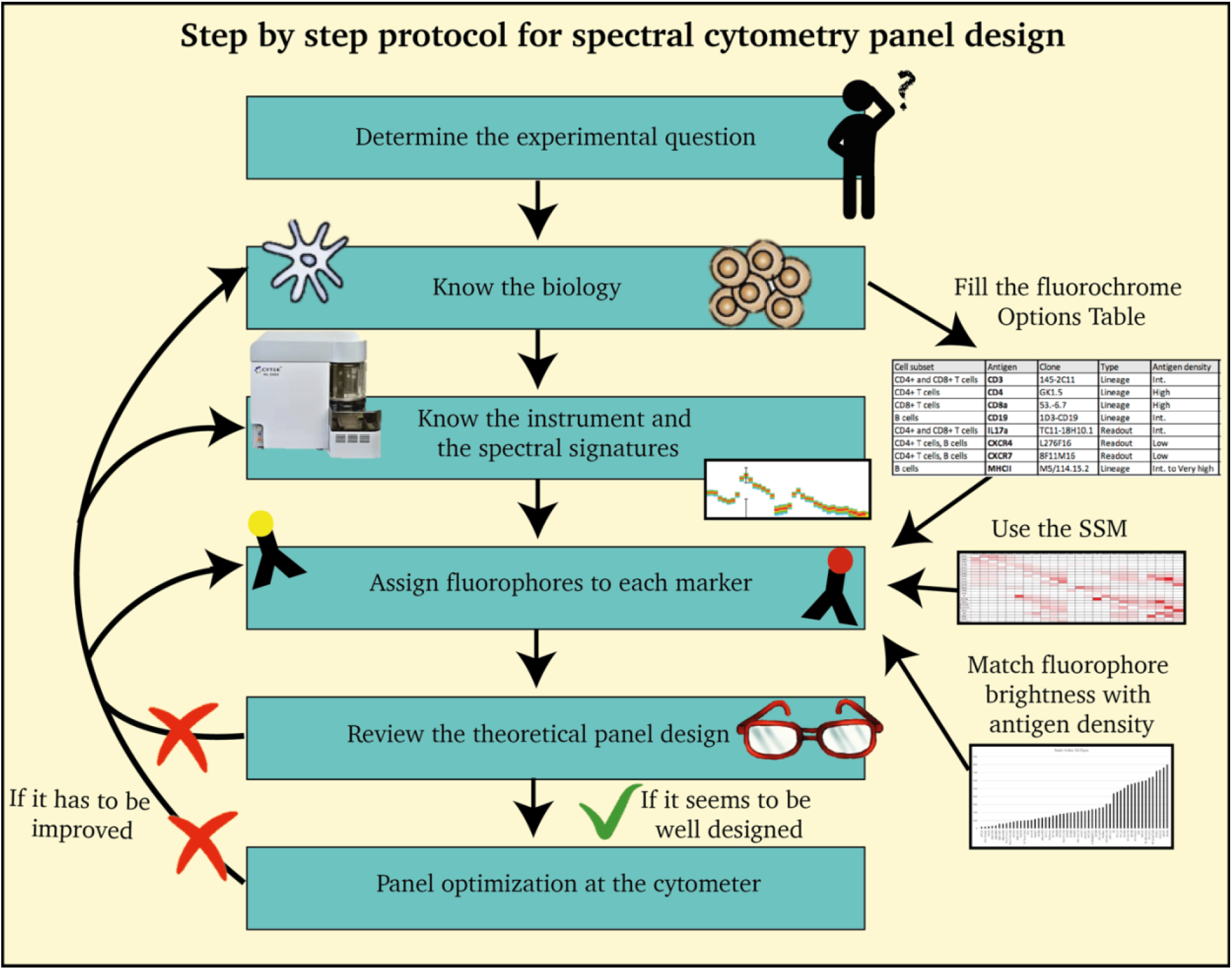
An overview of the step-by-step protocol that illustrates the design of a high-dimensional spectral flow cytometry panel.

### 1. Experimental question

The first step of every experiment, which also applies to spectral flow cytometric panel design, is to determine the objective and hypothesis of the study. Some questions you could ask are:

- What is the goal of the assay? Which questions are you willing to answer? *(for example, is IL-17A and CXCR4 expression in T cells increased during DSS colitis?, or such as, which immune cell types can be observed after temozolomide treatment of glioblastoma tumors?*
- Which readouts are needed to test your hypothesis? And which readouts are more critical than others? (*MFI, counts…*)
- Which tissues will have to be analyzed? What limitations are associated with the tissue? (*e.g. digestion, tissue stability, autofluorescence*)
- How will you need to process the samples to obtain the readout of interest? (*e.g. fixation, permeabilization, stimulation*)

### 2. Know the biology

Knowing the biology is the most critical part of panel design. It is imperative to know which markers are necessary to accurately define the cell types of interest and which markers are important for the functional readout necessary to address the experimental question.

To facilitate the collection of this information we suggest a table (an example can be found in the anticipated results, Table 2) to:

- Record the cell populations that need to be analysed.
- Define the lineage markers that are required to identify the cells of interest.
- Highlight co-expressed markers.

**Table 1.**
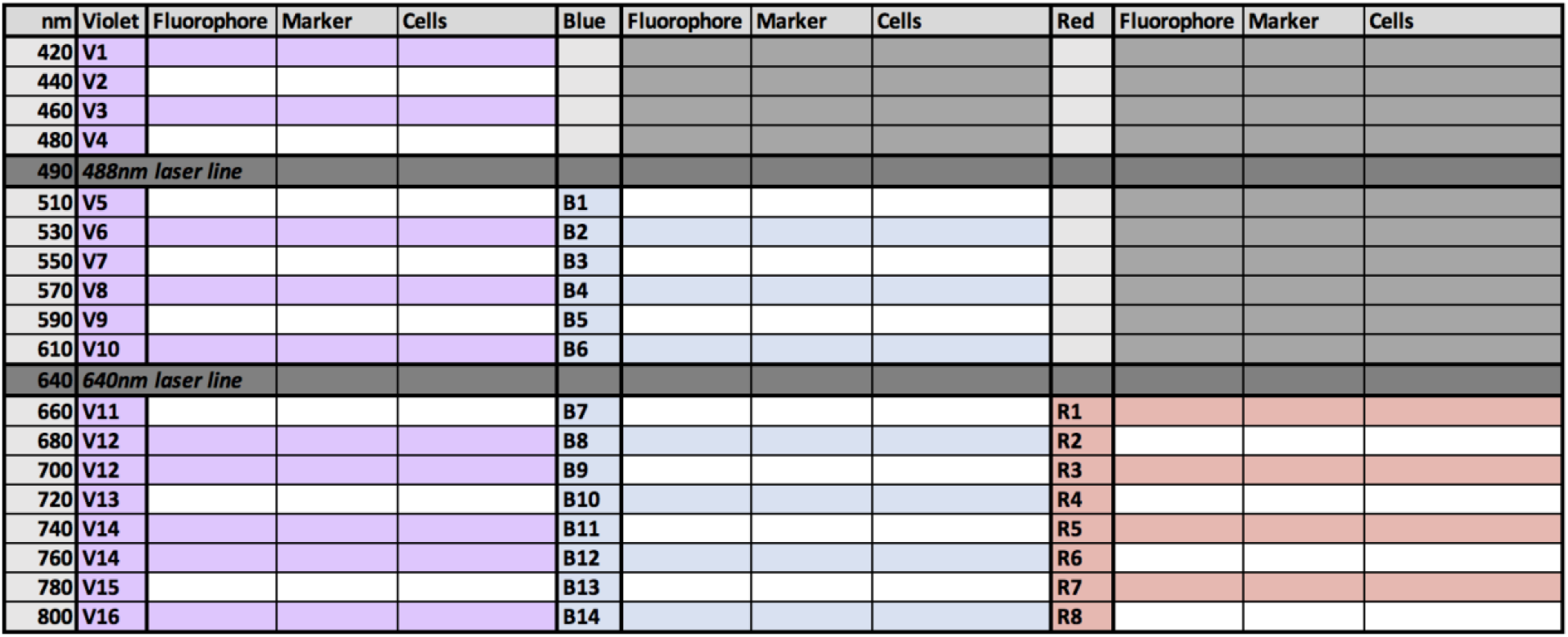
Panel Table. This table can be used to predict the spread induced on co-expressed markers. As a general rule, the closer the emission peak of two fluorophores are, the more spreading error and spillover will be observed between them. The table is structured to represent all the detectors of an Aurora 3L based on their wavelength range detection upon excitation by one of the 3 lasers. Each Marker – Fluorophore should be assigned on the row corresponding to the emission peak of the fluorophore. The main cell types expressing the marker should be annotated. Co-expressed markers should never be close in the same column, in order to avoid the emergence of spreading error. If possible, they should not be in the same row, as dyes showing a broad excitation spectrum will tend to emit in the same wavelength after being excited by different lasers.

| nm | Violet | Fluorophore | Marker | Cells | Blue | Fluorophore | Marker | Cells | Red | Fluorophore | Marker | Cells |
| --- | --- | --- | --- | --- | --- | --- | --- | --- | --- | --- | --- | --- |
| 420 | V1 |  |  |  |  |  |  |  |  |  |  |  |
| 440 | V2 |  |  |  |  |  |  |  |  |  |  |  |
| 460 | V3 |  |  |  |  |  |  |  |  |  |  |  |
| 480 | V4 |  |  |  |  |  |  |  |  |  |  |  |
| 490 | 488nm laser line |  |  |  |  |  |  |  |  |  |  |  |
| 510 | V5 |  |  |  | B1 |  |  |  |  |  |  |  |
| 530 | V6 |  |  |  | B2 |  |  |  |  |  |  |  |
| 550 | V7 |  |  |  | B3 |  |  |  |  |  |  |  |
| 570 | V8 |  |  |  | B4 |  |  |  |  |  |  |  |
| 590 | V9 |  |  |  | B5 |  |  |  |  |  |  |  |
| 610 | V10 |  |  |  | B6 |  |  |  |  |  |  |  |
| 640 | 640nm laser line |  |  |  |  |  |  |  |  |  |  |  |
| 660 | V11 |  |  |  | B7 |  |  |  | R1 |  |  |  |
| 680 | V12 |  |  |  | B8 |  |  |  | R2 |  |  |  |
| 700 | V12 |  |  |  | B9 |  |  |  | R3 |  |  |  |
| 720 | V13 |  |  |  | B10 |  |  |  | R4 |  |  |  |
| 740 | V14 |  |  |  | B11 |  |  |  | R5 |  |  |  |
| 760 | V14 |  |  |  | B12 |  |  |  | R6 |  |  |  |
| 780 | V15 |  |  |  | B13 |  |  |  | R7 |  |  |  |
| 800 | V16 |  |  |  | B14 |  |  |  | R8 |  |  |  |

**Table 2.**
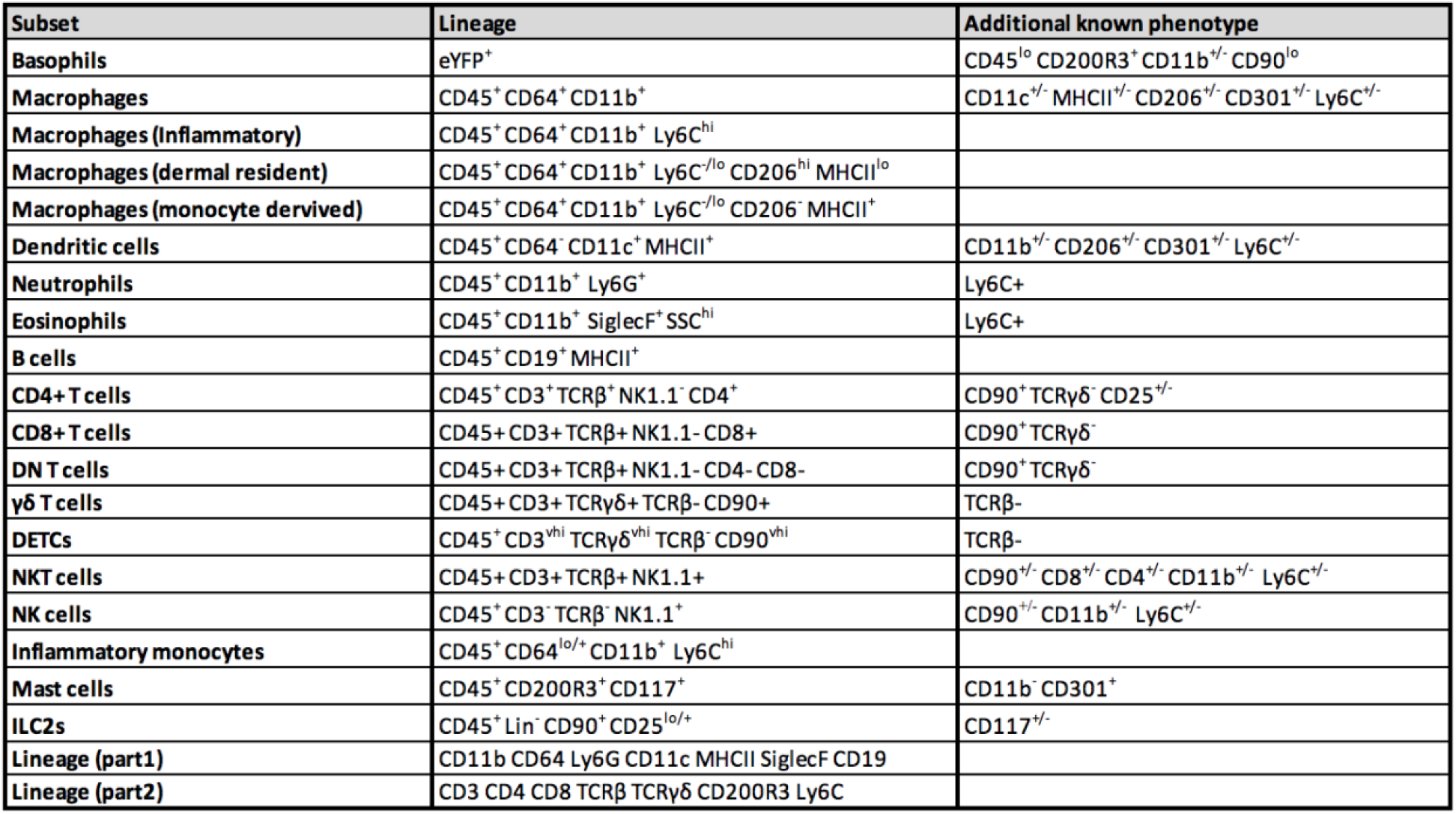
Cell Subset Table. Cell subsets, lineage markers and co-expression phenotype are listed.

The cell subsets table (table 2) allows us to visualize all the antigens of interest for the new panel. The antigens should then be listed in a second table, called antigen table (table 3). Columns should be filled by following these steps:

- Define if the markers are to be used as a readout or are lineage defining.
- Decide on the clone to be used (based on previous experience or publications).
- Note the expected antigen density (low, medium and high), if known.
- Record the availability of fluorophore-conjugated antibodies for each marker of interest. Commercial websites as well as specialized search engines such as biocompare.com or fluorofinder.com are useful tools to interrogate. If a certain clone is only available in a few fluorochromes, list these fluorochromes in the table. Otherwise note common and add only unusual fluorochromes (ex. APC-Cy5.5 or BV750).

**Table 3.** Antigens Table. Antigens, readout types, clones, antigen density and fluorophore availability are listed.

| Antigen | Type | Clone | Antigen density | Fluorophores available |
| --- | --- | --- | --- | --- |
| IL4 | Readout | Reporter | Low to high | AmCyan |
| IL13 | Readout | Reporter | Low to high | dsRed |
| MCPT8 | Lineage | Reporter | High | YFP |
| CD45 | Lineage | 30F11 | High | Common, APC-Cy5.5 |
| CD64 | Lineage | X54-5/7.1 | Low | Common |
| CD11b | Lineage | M1/70 | Low to very high | Common, AF594, BV570, BV750 |
| Ly6C | Lineage | HK1.4 | High | Common, BV570 |
| CD206 | Lineage | C068C2 | High | Common, AF594 |
| CD301 | Lineage | LOM/14 | Medium | Common, AF594 |
| CD11c | Lineage | N418 | Low | Common, AF594, BV570, BV750 |
| MHCII | Lineage | M5/114.15.2 | Very high | Common, AF594, BV570, BV750 |
| Ly6G | Lineage | 1A8 | High | Common, AF594, BV570 |
| SiglecF | Lineage | S17007L | Medium | APC, BV421, FITC, PE, SB436, PE-Dazzle594 |
| CD3e | Lineage | 145-2C11 | Low to very high | Common, PE-Cy5 |
| TCRb | Lineage | H57-597 | Medium | Common, AF594, BV570, PE-Cy5 |
| CD4 | Lineage | RM4-5 | High | Common, BV570, BV750, PE-Cy5 |
| CD8a | Lineage | 53-6.7 | High | Common, BV570, BV750, PE-Cy5 |
| TCRgd | Lineage | GL3 | High | Common |
| CD90.2 | Lineage | 53-2.1 | High | Common, BV570, PE-Cy5 |
| NK1.1 | Lineage | PK136 | Medium | Common, BV570, PE-Cy5 |
| CD19 | Lineage | 6D5 | Medium | Common, AF594, BV570, PE-Cy5 |
| CD200R3 | Lineage | Ba13 | Medium | APC, PE, PerCPCy5.5 |
| CD117 | Lineage | 2B8 | High | Common, PE-Cy5, BB700 |
| CD25 | Lineage | PC61 | Low | Common, BB515, AF594, PE-Cy5 |

### 3. Define fluorophores

Only dyes with unique spectral signatures should be used in the same panel. However, dyes with similar spectral signatures can be used at the risk of introducing spreading error, which can be mitigated if they are used with markers that are not co-expressed. The spectral signatures of many fluorophores have been extensively defined by Cytek Biosciences for their 1, 2, 3, 4 and 5 laser Auroras (https://spectrum.cytekbio.com/) and spectral signatures of additional fluorophores can be assessed using other online fluorescence spectra viewers (e.g. BD, Biolegend, Thermofisher, Expedeon), or by manually assessing the signature by acquiring single stained controls. Furthermore, the brightness of the fluorophore should also be considered.

### 4. Assign fluorophores to each marker

The most difficult task in multicolor panel design is to match the most appropriate fluorophore with each marker of interest. Fluorophore brightness and availability, levels of marker expression and marker co-expression, as well as spectral spillover have to be considered. Always match the fluorophore brightness with antigen expression levels by combining brighter fluorophores with weaker expression levels, and vice versa. The summary of stain indexes shown in Figure 2A provides a useful tool for fluorophore classification. Co-expressed markers and markers selected for functional readouts should be matched with fluorophores that receive minimal spread from other fluorophores in the panel. An indication of the spreading error between fluorophores can be found on the Cross Stain Index (CSI) matrix in Figure 2B. Determining spreading error (SE) is crucial when building high-dimensional fluorescence panels. Factor that contribute to the amount of SE from a given fluorophore into another fluorescence detector have been well described^15,16^. It is important to note that the CSI matrix is highly specific to the instrument setup and the fluorophores used and needs to be modified if additional or alternative fluorophores or another instrument are used.

**Figure 2.**
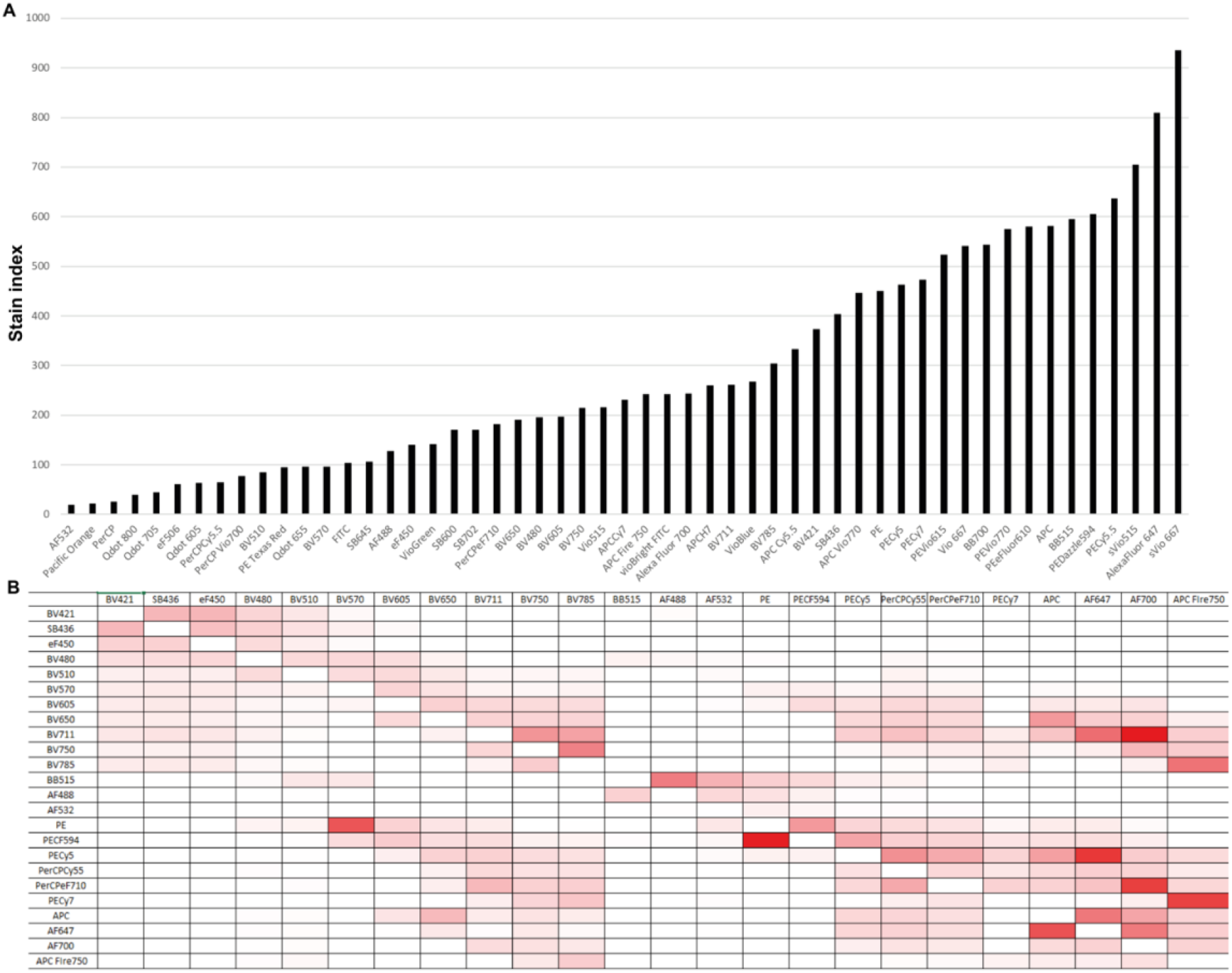
***(A)** Stain index of 53 commonly available dyes for the 3-laser Aurora (Cytek). Higher stain index represents brighter fluorophores or lower backgrounds. **(B)*** Cross Stain Index (CSI) matrix *for 24 unique-signature fluorophores that can be used in combination on a 3-laser Aurora spectral flow cytometer. Spillover is contributed by fluorophores listed in the rows spilling into the fluorophores listed in the columns (for example PE-CF594 spreads strongly into PE). Combinations depicted in red should be assigned to non-co-expressed markers or used for dump and viability channels. Note: the Stain index and the CSI matrix will vary depending on the instrument and laser configuration and are shown here for the 3-laser Aurora. (Cytek, also found online in Cytek website)*.

In order to facilitate the selection process, we suggest filling a Panel Table (see Table 1) using the following strategy:

- Note fluorescent reporter genes first, as they have a predefined fluorescent signature.
- Assign very rare markers to the only fluorophore options available (e.g. CXCR5 to PE).
- Assign the most common markers to rare fluorophores that could fit a distinct spectral signature in the panel (e.g. *CD4 to BV750*).
- Assign the remaining markers to other fluorophores considering their co-expression and spread.
- Unless non-viable cells are to be interrogated further, the viability dye can be selected last, regardless of any spreading issues due to the wide range of available fluorophores.

### 5. Review the theoretical panel design

In order to theoretically validate the panel before proceeding to practical testing, it is advised to review the panel on a marker-by-marker and cell-by-cell basis with the help of the annotated tables by asking the following questions:

- Are fluorophores that induce high amounts of spread allocated to non-co-expressed markers?
- Do the “readout” markers receive a minimal amount of spread?
- Are fluorophore brightness and antigen expression levels well matched?

To address potential issues, markers that are available in multiple fluorophores can be swapped out to see if spillover can be reduced. Additionally, fluorophores that create (but do not receive) the most spillover can be designated to dump or viability channels.

## COMMENTARY

Panel design considerations for a spectral cytometer overlap heavily with good standard practices for panel design on a conventional cytometer (i.e. ranking antigens and matching fluorophore brightness with antigen expression levels). However, spectral cytometry provides greater flexibility in fluorophore selection, as well as additional tools to help with successful multiparameter panel design, which are further outlined below.

### a) Instrument set up

On the Aurora, the “Cytek Assay Settings” are designed to provide optimized gains for all detectors. This optimization has been done in order to provide the best resolution of most known fluorophores on antibody stained cells. These gains are automatically adjusted after each daily QC based on laser and detector performance towards an ideal value, which is defined by the detection and optimal placement of proprietary fluorescent beads against predetermined target values. Due to the use of APDs, output fluorescence intensity increases linearly with gain up to the maximum value of 4×10^6^.

If the signal observed during testing is offscale using the optimized “Cytek Assay Settings”, it is generally better to titrate down the antibody used than to modify the instrument settings. However for special circumstances (such as fluorescent reporter gene expression or when saturating antibody titers are needed), the gains can be adjusted to allow dim or bright signals to be detected accurately. It is recommended to adjust the gains by a fixed percentage for all the detectors of a specific laser (SpectroFlo™ v2.0 and above), instead of adjusting the gains of individual detectors. It is extremely important to note that changing the gains of any detector will alter the spectral profile of all the fluorophores. This generally implies that the differences in spectral signatures between fluorophores will decrease, compared to the optimized default gain setting. Therefore, adjusting the gains away from the optimized default setting to enable detection of a specialized fluorophore might increase overall spreading error between other fluorophores, and should only be done if absolutely necessary. Single stain controls and all multi-stained samples will need to be acquired using the same settings, be it the “Cytek Assay Settings” or an adjusted version.

### b) Using spectral signatures

Fluorophores emit light over a range of wavelengths, and in conventional flow cytometry optical filters are used to capture peak fluorescence emission in a primary detector. When the emission profiles of two or more fluorophores overlap, the light emitted from one fluorophore appears in a non-primary detector (a detector intended for another fluorophore). This is referred to as spillover. Single-stained controls must be acquired to calculate the amount of spillover into each of the non-primary detectors. In conventional flow cytometry, spillover can be corrected by using a mathematical calculation called compensation.

In spectral cytometry the full emission spectrum of each single stained sample can be used to determine the contribution of each fluorophore in a mixed sample, using spectral deconvolution (unmixing) algorithms in real-time or post acquisition^17^. Here, the key to differentiating various fluorophores is to have distinct patterns or signatures across the full spectrum. Because the system measures the full range of emission (not only peak emission), two dyes with a similar emission maximum but different spectral signatures, such as APC and AF647, can be distinguished from each other (Figure 3).

**Figure 3.**
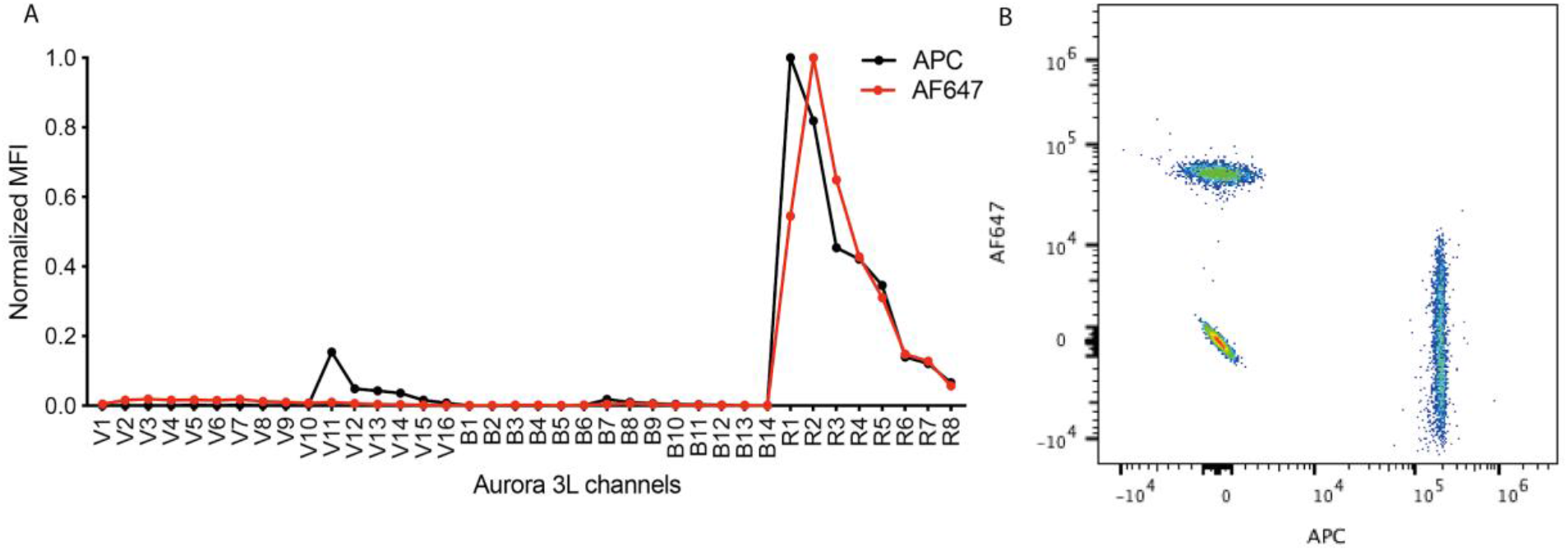
**(A)** Spectral pattern comparison (APC vs AF647); Note the stronger violet emission for APC compared with AF647 and differences in the peak emission profile **(B)** Dot plot showing that AF647 and APC stained beads can be discriminated by spectral flow cytometry, at the expense of significant spreading error.

### c) Fluorophore brightness and Stain index

The Stain Index (SI) is an equation to help determine the relative brightness of a fluorophore on a given instrument^16^ and can be calculated using the following formula:

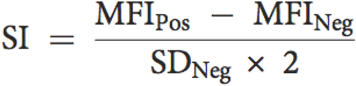

Absolute brightness depends on many attributes, namely the intrinsic fluorescence of the fluorophore, laser power, excitation wavelength, optical filters and detector type. Therefore, the SI is heavily dependent on the cytometer used and its specific setup.

An example of these differences is shown in Figure 4. We adjusted the mean fluorescence intensity (MFI) of common fluorophores on a conventional cytometer to match those measured the Aurora using optimized defaults (Figure 4A). This resulted in an increased background noise of the negative population on the conventional cytometer (measured by the robust Standard Deviation, rSD, Figure 4B). The rSD was smaller on the Aurora in most channels and generally allowed better SI values (Figure 4C). As previously reported^11^, APDs perform better at wavelengths above 650nm where the higher quantum efficiency contributes to better signal-to-noise ratios than PMTs. This was also reflected in our measurements resulting in a higher SI for dyes emitting in red and far red channels (Figure 4C).

**Figure 4.**
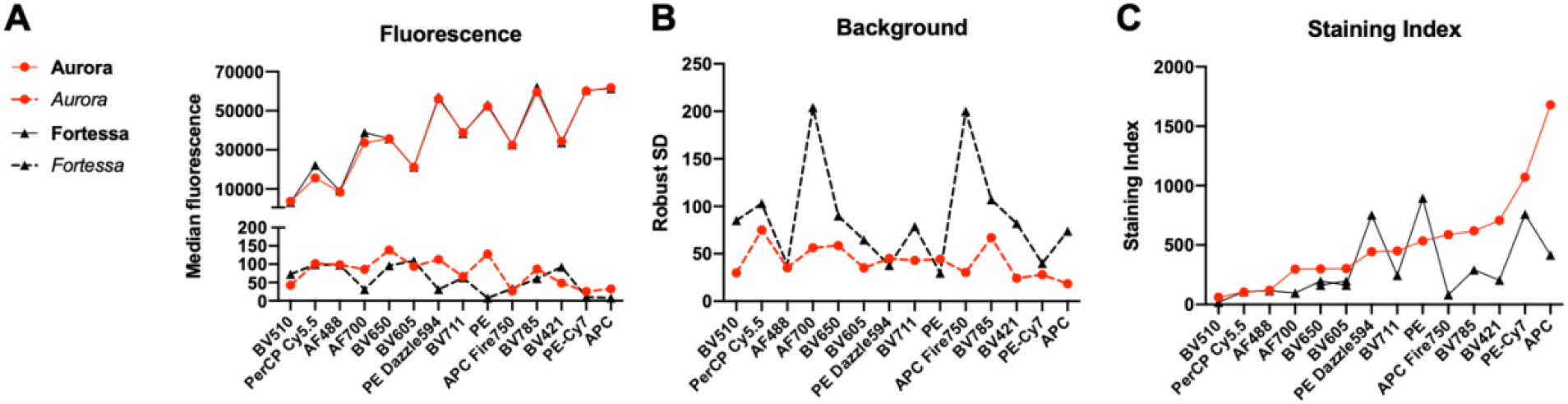
Brightness comparison of different fluorophores conjugated with anti-CD4 antibodies on human peripheral blood leukocytes. **(A)** Raw median fluorescence for antibody positive (solid line) and antibody negative (dashed line) populations **(B)** Background noise (standard deviation of the negative population) **(C)** Stain Index (SI). **(A, B, C)** Fluorophores are listed based on their increased Stain Index according to the Aurora data. The median fluorescence measured on the Fortessa has been adjusted to match those measured on the Aurora 3L using optimized defaults.

### d) Autofluorescence (AF)

AF is a natural characteristic of all cells whereby biological substances and structures within the cell fluoresce. AF can be attributed to biomolecules such as NADH, folic acid, and retinol which have emission maxima in the range of 450–500 nm, and riboflavin, flavin coenzymes, and flavoproteins which have emission maxima in the range of 520–540 nm.^18^. AF creates a background that can impair the detection of dim markers emitting light at the same wavelengths In spectral flow cytometry, the spectral profile of unstained cells can be collected and treated as an independent parameter (AF), which allows the AF signature to be extracted using the unmixing algorithm. Correcting for autofluorescent signatures can improve the signal-to-noise ratios (Figure 5) and enables a clearer distinction of fluorophores that have been avoided in the past, due to their peak emission in the AF range (e.g. AF532 and BV510). Furthermore, AF correction can improve the resolution of certain markers in highly autofluorescent tissues such as the brain, lung, skin, intestine and tumors^9^.

**Figure 5.**
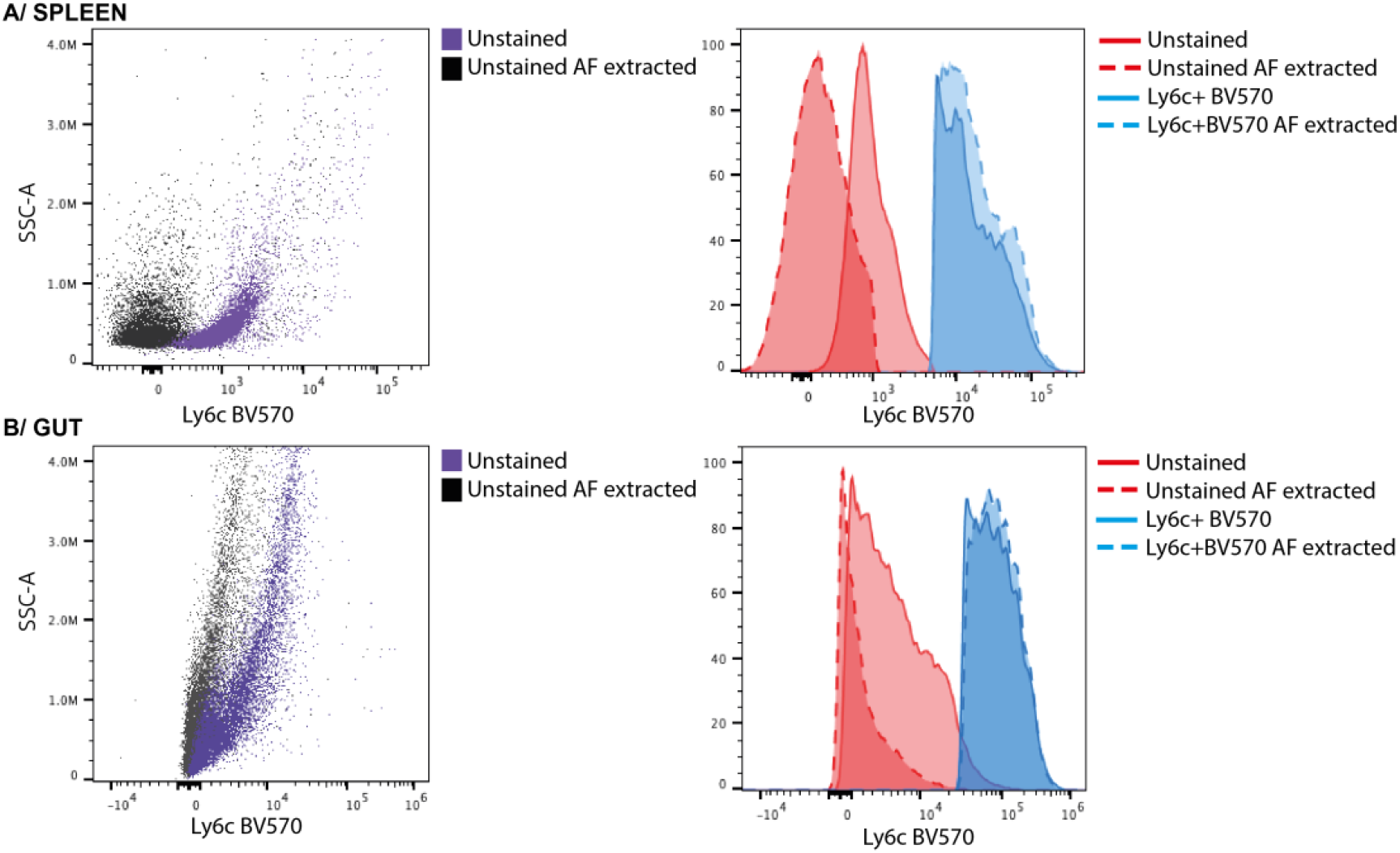
**Autofluorescence (AF) correction** in **(A)** spleen and **(B)** gut samples stained with anti-Ly6C BV570. The correction of AF improves positive signal resolution by lowering the background of the negative population with minimal effect on the positive signal.

## CRITICAL PARAMETERS

### a) Antibody titration

Antibody titration is an absolute requirement for the development of a successful high-dimensional spectral cytometry panel. As with conventional flow cytometry, optimal antibody titration is necessary to reduce the amount of antibody used, reduce background signal by minimizing non-specific binding, and to compare samples accurately^19^. If a signal is too bright, the best practice is to use an antibody conjugated to a dimmer fluorophore. If this is not possible, it is better to titrate an antibody down to reduce the spread created in other channels^20^, than to change the instrument settings. However, antibodies used to assess functional readouts should not be titrated below saturation, as saturating concentrations are needed to avoid variability and ensure accurate detection between samples^21^.

### b) Reporter genes

For accurate detection of fluorescent reporter gene expression, single stained controls should be derived from the reporter system under experimental conditions that induce the highest level of gene expression (e.g., stimulation), whereas all other single stains should be derived from wild type cells. Furthermore, fully stained wild type cells serve as the Fluorescence Minus One (FMO) control for the fully stained gene reporter samples. It is important to ensure that fluorescent reporter gene expression is on scale, otherwise the gains will need to be adjusted. However, modifying the gains of the “Cytek Assay Settings” will impact all channels resulting in the suboptimal detection of the remaining fluorophores. An extreme example is shown Figure 6, where dramatically reduced gains in the red and blue laser detectors resulted in a distortion of the spectral signature and a strong increase in spreading error.

**Figure 6.**
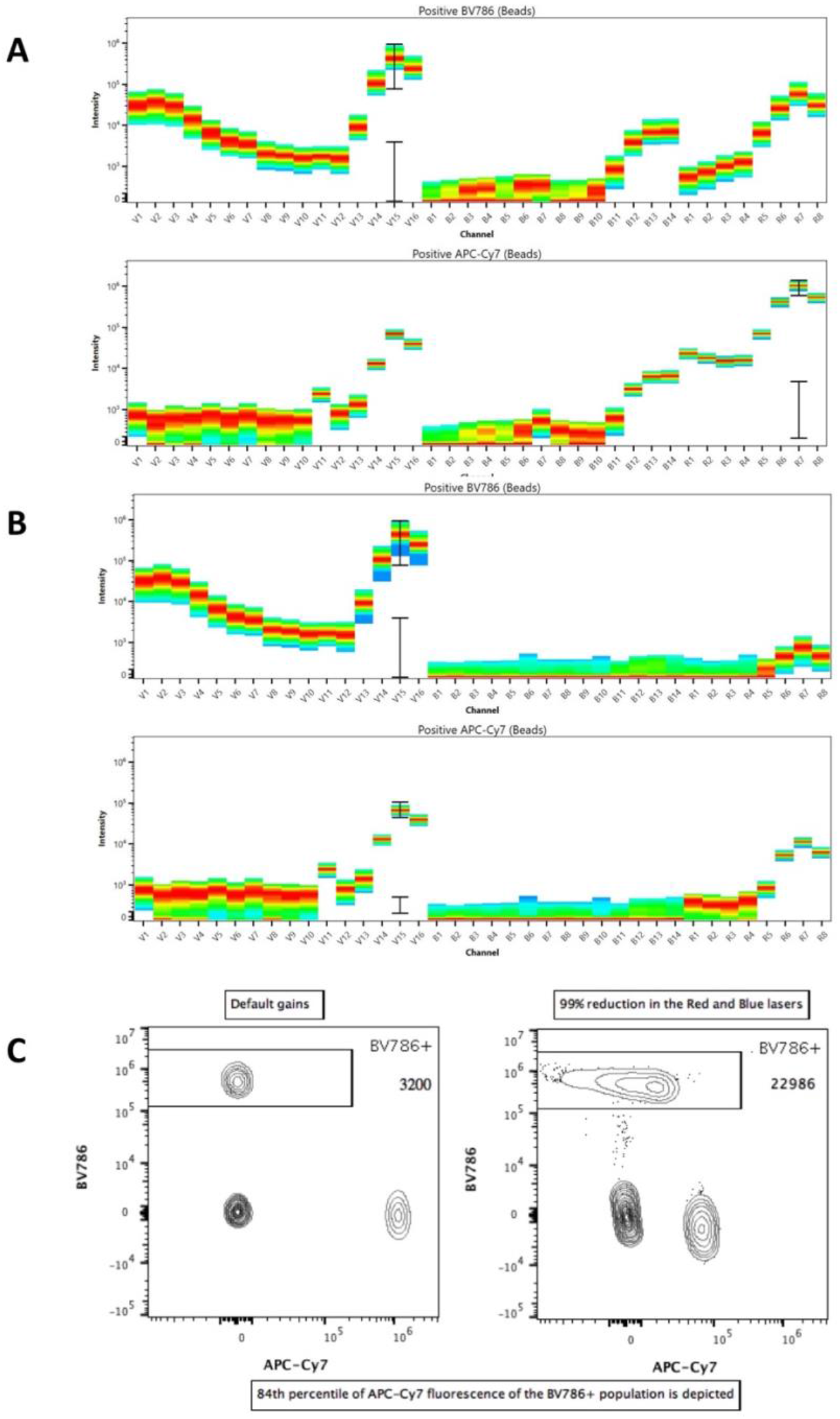
Compensation beads were stained with BV786 or APC-Cy7 conjugated antibodies and their spectral profile was acquired on an Aurora 3 laser system **(A)** using vendor established gains (“Cytek Assay Settings”) or **(B)** after a 99% reduction of the gains of all detectors associated with the red and blue lasers. Unmixing was performed for both data sets, and then single stained bead preparations were acquired in default or reduced gains conditions **(C).** The 84th percentile of the APC-Cy7 fluorescence detected on the BV786+ beads after spillover compensation (a measure of the SE^16^) is depicted on the dot plots. Note the width basis for data display needed to be further adjusted post gain reduction to produce the single BUV786+ population.

### c) Tandem Dyes

Many available fluorophores are tandem organic or polymer dyes, in which the excitation of the donor fluorophore leads to an emission by the acceptor dye of the tandem^22^. The resulting spectral profile is a hybrid between the excitation and emission spectra of both dyes. However, spectral profiles of tandem dyes slowly degrade over time, are different in a lot-to-lot fashion (due to changing acceptor/donor ratios), and can also be affected by experimental conditions, such as fixation and incubation at 37°C^22,23,24^. Therefore, single stained controls need to use the same antibody lot and be treated in the same manner as the experimental samples. New lots of tandem dyes need to be re-titrated to ensure consistent results.

### d) Spreading error

Spreading error (SE) is a large contributor to reduced signal resolution^16^. To minimize spreading error, only fluorophores with a unique spectral signature should be selected. While spectral flow cytometry can distinguish between dyes that have a very similar signature (Figure 3), using these fluorophores will introduce SE and should not be used to detect co-expressed markers. As in conventional flow cytometry, SE is a direct function of fluorescence intensity, and as such, panels should be designed so that fluorophores that contribute the most spread are paired with dimly-expressed markers.

### e) Dump channels

It is advisable to use fluorophores with identical spectral signatures for dump channels. Therefore, the use of tandem dyes should be avoided, with the exception of PerCP-Cy5.5 which is a stable tandem. Fluorophores that produce spreading error (e.g. PerCP-Cy5.5) or fluorophores that emit in the AF range (e.g. BV510, V500 or eFluor506) are ideal candidates for use as dump channel dyes, as the spreading error created by the positive signal will be excluded from further analysis. Spreading error from other fluorophores into the dump channel should be avoided, as cells of interest could accidentally be excluded. The same rationale can also be applied for viability dye selection.

### f) Spectral reference controls (SRC)

SRCs are single stained controls used for establishing reference vectors to enable successful linear unmixing to occur. SRCs should always be as bright or brighter than experimental samples, ideally with a fluorescence intensity above 5×10^5^ but below 4×10^6^, the maximum value of the linear range of detection for Aurora spectral flow cytometers. While compensation beads offer a convenient choice for SRCs, certain conditions, such as high antigen density^15^, might require the use of cells. It is recommended to use single stained cells of a similar tissue as the experimental sample as SRCs, if the positive signal observed is frequent and bright enough. It is essential to collect enough total events for the software to clearly distinguish the spectral fingerprint for both the positive and the negative populations.

SRCs should also be treated the same way as the fully stained samples, as fixation steps or pH variations can alter the spectral profiles of some fluorophores^15^. Such treatments can also affect the fluorescent signature of compensation beads, which should always be checked. One example is the use of BD Brilliant Staining Buffer, which is used to decrease the interactions of the Brilliant dyes among themselves and mitigate nonspecific signals in multicolor samples. However, this buffer also significantly alters the spectral profile of some latex beads (e.g. UltraComp beads, see Figure 7), and should be avoided when staining beads.

**Figure 7.**
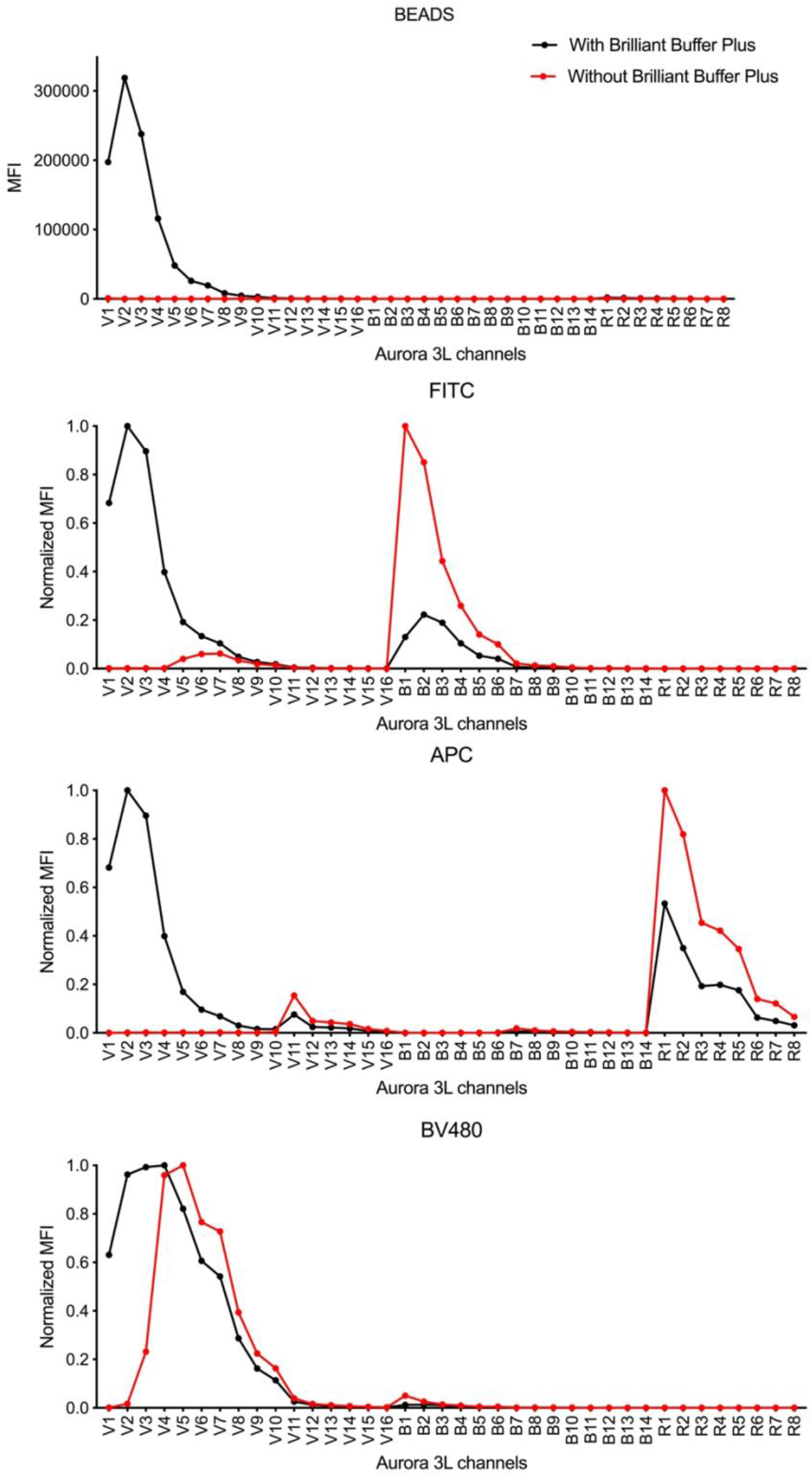
Ultracomp eBeads™ (ThermoFisher) were stained either with Brilliant stain buffer plus™ (BBP, BD Biosciences) or without (no BBP). Examples for beads and the fluorochromes FITC, BV480 and APC are shown.

Unstained SRCs have to be acquired for every experiment, and it is advised to use a matched unstained control for unmixing each individual tissue if multiple tissues are analysed, especially if AF correction is applied during the unmixing process.

The Aurora creates Raw and Unmixed FCS 3.1 files for each experiment. If an SRC did not yield and ideal signal or signature at the time of recording the experiment (e.g. not bright enough or included nonspecific signal such as autofluorescent cells), recorded data can be re-unmixed within the SpectroFlo™ software post acquisition using different SRCs. Similarly, samples can be re-unmixed with or without AF extraction.

Once optimal SRCs have been defined for a panel, they can be recorded for future use in the reference library within the software. When the controls are recorded within the setup and QC module, the data defining the controls is normalized to the detector gains and will be optimally adjusted during the daily setup process. Reference library SRCs should be re-acquired after any system stability or QC performance issue. It is important to note that new SRCs must be recorded for each new antibody lot, as there is significant lot-to-lot variation in fluorophore spectral profiles, particularly for tandem dyes.

## TROUBLESHOOTING

Panel optimization is an important step for any multicolor panel design. One of the first steps is to compare how antibodies behave in combination compared to single stained cells. By comparing the fluorescence intensity and spreading error of all single stained cells to the fully stained sample one can easily assess whether the combination of antibodies in the multicolor panel decreases the resolution of a given fluorophore by, for example, introducing spillover or spreading error. It is then easy to determine which fluorophore in the multicolour panel is reducing the resolution of the fluorophore of the single stain sample.

Not all FMOs will be required for the panel optimisation step. Only antibodies that may be creating spillover or spreading errors in a certain channel, will be acquired as FMOs to help with panel QC. FMOs or isotype controls are needed to reliably and reproducibly define the boundary between positive and negative readout signals across different cell populations and experiments. They are necessary for establishing gating boundaries between dimly expressed antigens and background and are an important tool for assessing panel performance.

If a conflicting pair of antibodies has been identified by assessing the single stained cells or FMO controls, alterations should be made to the panel by changing the antibody concentration or modifying the antibody-fluorophore assignment. Small corrections to spillover can be made by altering the compensation matrix within SpectroFlo™ or in a third-party software of choice.

## ANTICIPATED RESULTS

In this section we describe the design of a 25-parameter panel, which includes the use of triple reporter (YFP/AmCyan/DsRed) Basoph8×4C13R mice with or without experimental treatment. Each step of the basic protocol identified in Figure 1 is described and important considerations are highlighted.

### 1) Experimental question

The aim was to understand which cell types express IL-4 and IL-13 in the skin in a MC903 induced atopic dermatitis model using the triple reporter Basoph8×4C13R mice. These mice express eYFP as a specific marker for basophils, and report the expression of the type 2 cytokines IL-4 and IL-13 as AmCyan and dsRed, respectively. Experimental readouts were the expression of IL-4 (AmCyan) and IL-13 (dsRed) by the various immune cells identified. In order to analyse the fluorescence of these reporters, Basoph8xC57BL6 samples were used as control groups to define the IL-4 and IL-13 expression boundary on each cell population of interest.

Our analysis focused on the skin, a highly autofluorescent tissue requiring collagenase digestion to obtain single cell suspensions. Prior analysis of splenocytes has revealed that the expression of certain epitopes (i.e. cKit, CD3, CD4) was decreased by the digestion so bright fluorophores were needed to detect these markers. Furthermore, preliminary experiments revealed that reporter fluorophore stability was strongly affected by fixation and permeabilization, so only fresh cells were assessed.

### 2) Biology

Different subsets of T cells (including CD4+, CD8+, *γδ*, and dendritic epidermal T cells (DETC)), mast cells, basophils, ILC2s, eosinophils, NK cells, NKT cells, monocytes and neutrophils have been shown to express IL-4 and/or IL-13 during allergic inflammation^25–29^. To address whether these and other cell types expressed IL-4 or IL-13 in the skin after MC903 treatment we wanted to quantify these cell subsets and various macrophages subpopulations, as they are known to be responsive to type 2 cytokines. We listed the lineage defining phenotype of every cell subset of interest in a Cell subset Table (Table 4), and then listed all the antigens of choice in an Antigen table (Table 5) to annotate the associated cell subsets, clones, readout characteristics, antigen density and fluorophore availability.

**Table 4.** Example of a 25-color panel to detect type 2 cytokine expression of skin immune cells. The table was first filled with reporters (red), then essential or rare markers (orange), followed by different lineage markers highlighted in yellow, blue, and white.

| nm | Violet | Dye | Marker | Cells | Dilution | Blue | Dye | Marker | Cells | Dilution | Dye | Fluor. | Marker | Cells | Dilution |
| --- | --- | --- | --- | --- | --- | --- | --- | --- | --- | --- | --- | --- | --- | --- | --- |
| 420 | V1 | BV421 | SiglecF | Eosinophils | 100 |  |  |  |  |  |  |  |  |  |  |
| 440 | V2 |  |  |  |  |  |  |  |  |  |  |  |  |  |  |
| 460 | V3 | Pacific Blue | MHCII | DCs MACs | 10000 |  |  |  |  |  |  |  |  |  |  |
| 480 | V4 |  |  |  |  |  |  |  |  |  |  |  |  |  |  |
| 490 | 488nm laser line |  |  |  |  |  |  |  |  |  |  |  |  |  |  |
| 510 | V5 | AmCyan | IL4 | Basophils, T cells | Reporter | B1 | BB515 | CD25 | Lymphoids | 100 |  |  |  |  |  |
| 530 | V6 |  |  |  |  | B2 | YFP | YFP | Basophils | Reporter |  |  |  |  |  |
| 550 | V7 | BV510 | CD8a | CD8+ T cells | 200 | B3 |  |  |  |  |  |  |  |  |  |
| 570 | V8 | BV570 | CD11b | Myeloids | 100 | B4 | PE | CD301 | M2 / DCs | 100 |  |  |  |  |  |
| 590 | V9 |  |  |  |  | B5 | dsRed | IL13 | ILCs T cells | Reporter |  |  |  |  |  |
| 610 | V10 | BV605 | TCRb | CD4+ / CD8+ T cells | 100 | B6 | PE-Dazzle 594 | CD64 | MACs | 100 |  |  |  |  |  |
| 640 | 640nm laser line |  |  |  |  |  |  |  |  |  |  |  |  |  |  |
| 660 | V11 | BV650 | NK1.1 | NKs | 100 | B7 |  |  |  |  | R1 | AF647 | CD206 | M2 / DCs | 800 |
| 680 | V12 |  |  |  |  | B8 | PE-Cy5 | CD3 | T cells | 600 | R2 | APC | CD200R3 | Basophils Mast cells | 100 |
| 700 | V12 |  |  |  |  | B9 | BB700 | cKit | Mast cells | 200 | R3 | APC-Cy55 | CD45 | all | 50 |
| 720 | V13 | BV711 | Ly6G | Neutrophils | 200 | B10 | PerCPeF710 | TCRgd | gd T cells | 2000 | R4 | AF700 | Ly6C | Monocytes | 400 |
| 740 | V14 | BV750 | CD4 | CD4+ T cells | 100 | B11 |  |  |  |  | R5 |  |  |  |  |
| 780 | V15 | BV786 | CD11c | DCs MACs | 100 | B12 |  |  |  |  | R6 | Zombie NIR | Dead | Dead | 2000 in PBS |
| 800 | V16 |  |  |  |  | B14 | PE-Cy7 | CD90.2 | Lymphoids | 2000 | R8 | APC-CH7 | CD19 | B cells | 100 |

### 3) Know the instrument and the spectral signatures

Cytek 3L Aurora was used for this experiment. By using online fluorescence spectra viewers, we were able to identify 25 dyes with distinct signatures that could be used in our panel.

### 4) Assign markers to fluorophores

We sequentially assigned fluorophores to our markers of interests in a Panel table (such as Table 1). An overview of the panel design is illustrated in Table 4, where the main steps of the fluorophore assignment sequence have been colour coded.

Our primary readouts of interest were the IL-4 (AmCyan) and IL-13 (dsRed) expression by different cell types. All of our cell types of interest expressed CD45 as they were leukocytes, and it was therefore important to have a bright CD45 signal that did not spread in our readout channels. We chose APC-Cy5.5 because of its unique spectral signature and the rarity of this dye on the market. Similarly, we expected CD4+ CD3+ Th2 cells to be a major source of both IL-4 and IL-13. CD4 expression is quite selective and relatively high, so we assigned it to a less common fluorophore that creates and receives some spread, such as BV750. CD3 expression is very heterogeneous in the skin: αβT cells show low expression after digestion, while DETC show a very high expression. We needed a dye bright enough to be able to detect the CD3^low^ population, and to gate on CD3^-^ cells, while keeping CD3^hi^ cells on scale. Here we chose PE-Cy5, which is moderately bright on the Aurora 3L.

The commercial availability of CD200R3 antibodies is sparse and we were limited to APC and PE. As basophils and mast cells express CD200R3 and can express IL-13, we chose APC for our panel so we could not create any spread in the IL-13 dsRed channel. We knew from the literature that cKit is strongly expressed by mast cells^30^, but preliminary testing showed that its expression was decreased upon digestion. We therefore assigned it to a bright marker (BB700) in order to have a resilient identification of mast cells in our panel.

CD90.2 is strongly expressed in lymphoid cells. However, as the main positive lineage marker for ILC2s we needed to have a very bright fluorophore without much spread from other channels, and chose to assign it to PE-Cy7. Other T cell markers are well defined and quasi-exclusive and could be assigned later, away from AmCyan or dsRed.

We also wanted to analyse the different macrophage subsets in the skin, which are not known to express type 2 cytokines. The expression of CD64 can be dim, so we chose to assign it to Pe-Dazzle594, a very bright dye. Macrophages also express CD11b at high levels, but it is also expressed by many other cell types at lower levels. However, only the high expression of CD11b is lineage-defining in our panel, so we chose to assign it to the dim BV570. Macrophages in the dermis are also known to express CD11c, which is a key lineage marker for dendritic cells. We assigned it to a bright dye (BV786) far from any interference with other macrophage markers.

Similarly, CD206 and CD301 expression define type 2 subsets of macrophages and dendritic cells in the skin, but are not expressed by other cells. We assigned them to AF647 and PE, based on availability. MHCII expression is critical to identify dendritic cells, and is highly expressed. We assigned it to Pacific Blue, a dim marker. Ly6C expression is very high on inflammatory macrophages and monocytes, so it was assigned to AF700, far from other macrophage or monocyte markers.

More general markers, such as lineage defining markers for eosinophils, neutrophils, NK cells, and B cells (SiglecF, Ly6G, NK1.1, CD19) were assigned to remaining available fluorophores (BV421, BV711, BV650 and APC-H7, respectively). As these cell types could potentially express IL-4 and IL-13 they were assigned to dyes that were spectrally different from AmCyan and dsRed.

Other T cell markers were then assigned based on availability (TCR*β* BV605, TCR*γδ* PerCP-eF710 and CD8 BV510). Among one of the last markers war CD25, which is dimly expressed on T cells and was thus assigned to BB515. Finally, the viability dye was assigned to Zombie NIR, based on the few free channels still available.

### 5) Panel review

Several pairs of dyes used in this panel are impacted by significant spreading error between them (such as AF647/APC and BB515/YFP. However, this effect was mitigated by assigning these fluorophores to non-co-expressed markers (e.g., CD206 and CD200R3, for AF647 and APC). Other difficult combinations included PE/dsRed/PE-Dazzle594, PE-Cy5/BB700/PerCP-eF710 and BV605/BV650/PE-Cy5, as these combinations are known to be co-expressed on certain cell types. We took note of these particular fluorophore combinations and investigated their separation during the panel testing phase.

### 6) Panel Testing and optimization

Testing the panel on the unstimulated ear skin revealed that all the populations could be resolved adequately (see Figure 9). However, during experimental inflammation of the skin, IL-4 expression of some macrophages was detected, creating spreading errors between IL-4 (AmCyan) and MHCII (Pacific Blue). We therefore reduced the concentration of MHCII-Pacific Blue from 1/2000 to 1/10000. This change considerably improved the resolution of IL-4 expression on macrophages, without impairing our ability to detect other MHCII+ cell types such as dendritic cells (Figure 8).

**Figure 8.**
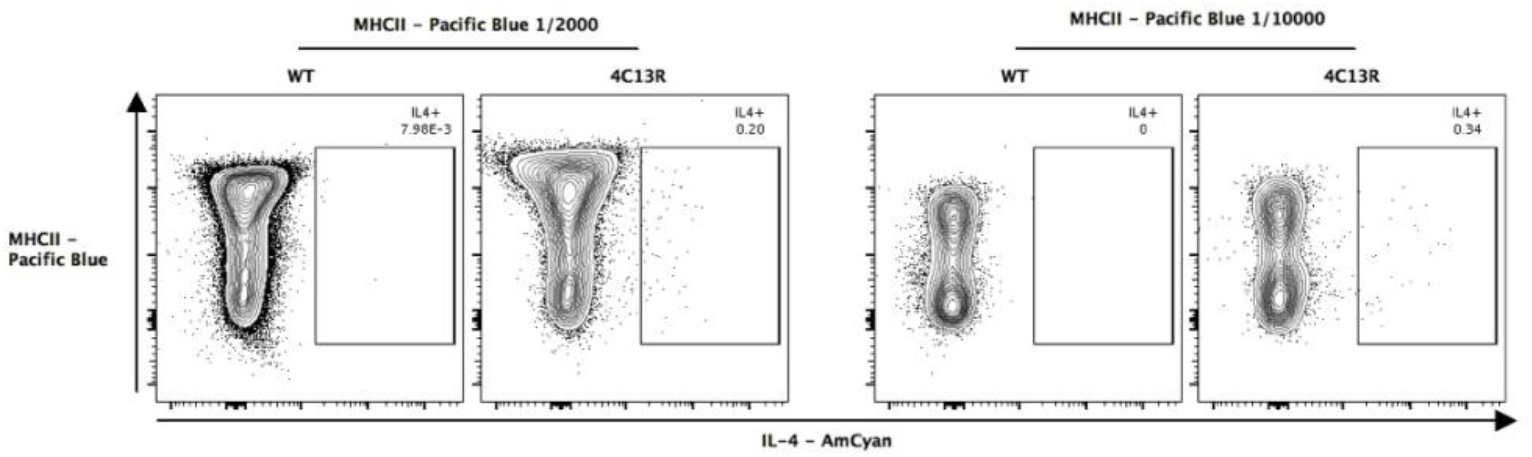
The spreading error of Pacific Blue MHCII in the AmCyan channel. Analysis of MC903 treated wild type (WT) ear skin revealed that the high expression of MHCII by CD45+ CD64+ CD11b+ macrophages was impeding the detection of IL4-AmCyan in the reporter mouse. Reducing the concentration of the MHCII-Pacific Blue antibody from the 1/2000 (left panel) to the 1/10000 dilution (right panel) did improve the detection of IL-4 by reducing the spreading error of Pacific Blue into the AmCyan channel.

**Figure 9.**
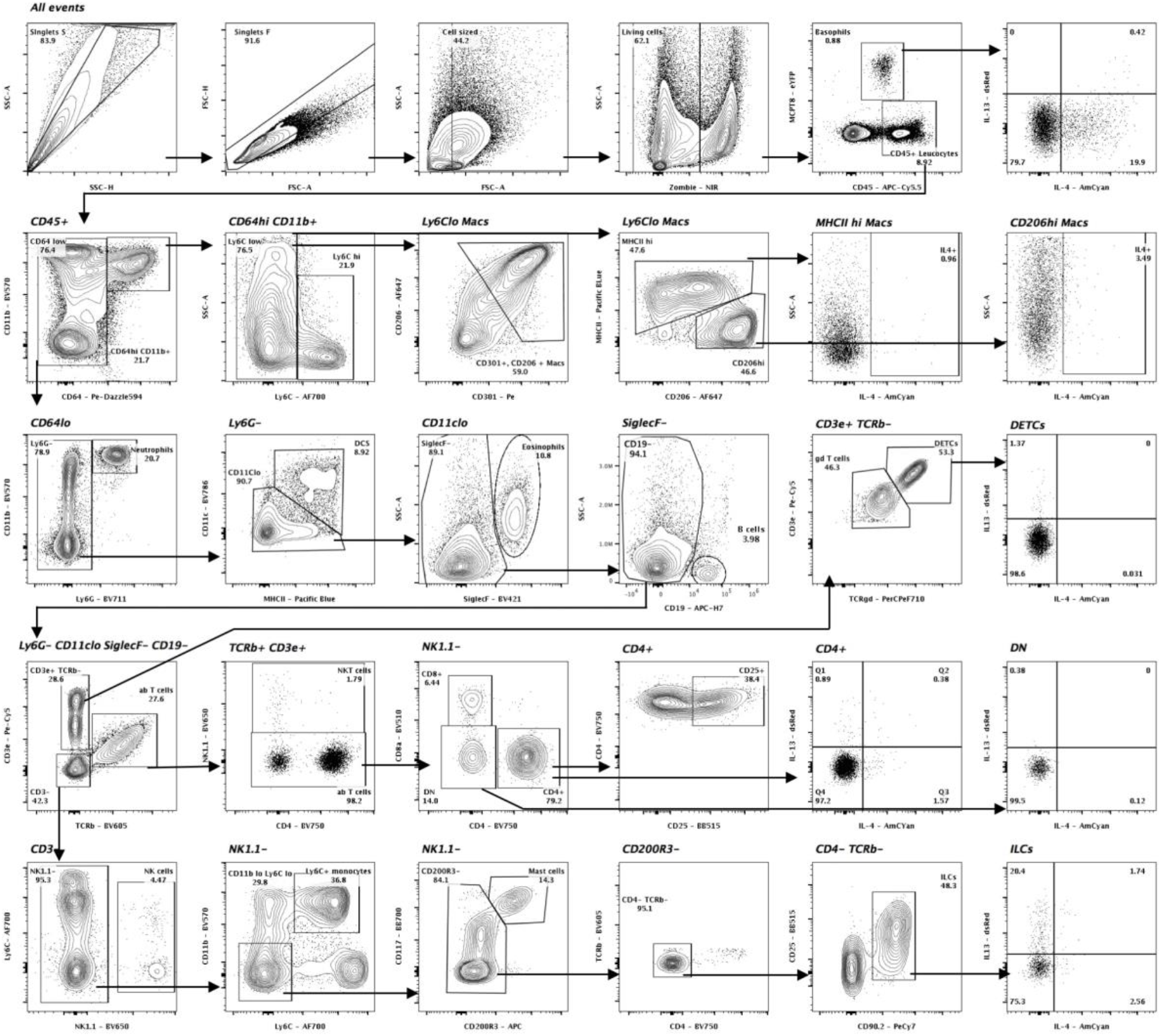
Gating strategy of the full 25 parameter staining panel of MC903 treated ear skin represented as contour plots gated in cascade. CD45+ leucocytes and YFP+ basophils are gated from live single cells. CD64+ CD11b+ macrophages were comprised of a Ly6C+ inflammatory subset and two Ly6^lo^ subsets, MHCII^hi^ or CD206^hi^. From the CD64^lo^ population, Neutrophils are observed as CD11b+Ly6G+. In the Ly6G-population, dendritic cells were defined as MHCII+ CD11c+. Eosinophils were gated as SiglecF+ SSC^hi^ cells from the CD11c^lo^ subset, as were CD19+ B cells. Within the Ly6G-CD11c^lo^ Siglec F-CD19-population, TCRβ+ CD3+ T cells and CD3+ TCRβ-T cells were identified. αβT cells contained NK1.1+ NKTs, CD8+, CD4+ and CD4-CD8-Double Negative (DN) T cells. The CD4+ subset further contained a population of CD25+ T cells. The TCRβ-population could be divided in two subsets of γδT cells, including a CD3/CD90.2/γδTCR^vhi^ Dendritic epidermal T cell population (DETC). CD3 negative subsets were gated as NK1.1+ NK cells, CD11b+ Ly6C+ inflammatory monocytes, CD11b-Ly6C-CD117+ CD200R3+ Mast cells, or Lineage–CD90.2+ CD25+ ILC2s. The expression of IL-4 and IL-13 by relevant immune cells was also analysed.

### Time considerations

Designing, validating and analysing high-dimensional spectral flow cytometry panels takes significantly more time than running the actual experiment. It can take between 2 weeks and 1 month to design a well optimized panel. It is therefore advisable to develop certain key panels that can be applied to several experimental questions and models. The time required for the panel design and optimization will also depend on the amount of time required between ordering the antibodies and receiving them.

Week 1: Panel design and antibody ordering.

Week 2: Antibody titrations and acquisition of Single stained cells and FMOs (if needed). Panel review.

Week 3: Second or third iteration of the panel if changes in fluorophore(s) or titre(s) have to be made.

## Acknowledgements

This work was enabled by the Hugh Green Cytometry Centre and we wish to thank the Hugh Green Foundation for their ongoing generosity and support. The authors are also very grateful to Kathy Muirhead, Katherine Woods, Joanne Lannigan and Janelle Shook for their expert opinions and editing prowess.

